# Reliable analysis of clinical tumor-only whole exome sequencing data

**DOI:** 10.1101/552711

**Authors:** Sehyun Oh, Ludwig Geistlinger, Marcel Ramos, Martin Morgan, Levi Waldron, Markus Riester

**Affiliations:** Graduate School of Public Health and Health Policy, City University of New York, 55 W 125th St, New York, NY 10027, USA.; Institute for Implementation Science and Population Health, City University of New York, 55 W 125th St, New York, NY 10027, USA.; Roswell Park Cancer Institute, 665 Elm St, Buffalo, NY 14203, USA.; Novartis Institutes for BioMedical Research, 250 Massachusetts Ave, Cambridge, MA 02139, USA.

## Abstract

**Background:** Allele-specific copy number alteration (CNA) analysis is essential to study the functional impact of single nucleotide variants (SNV) and the process of tumorigenesis. Most commonly used tools in the field rely on high quality genome-wide data with matched normal profiles, limiting their applicability in clinical settings.

**Methods:** We propose a workflow, based on the open-source PureCN R/Bioconductor package in conjunction with widely used variant-calling and copy number segmentation algorithms, for allele-specific CNA analysis from whole exome sequencing (WES) without matched normals. We use The Cancer Genome Atlas (TCGA) ovarian carcinoma (OV) and lung adenocarcinoma (LUAD) datasets to benchmark its performance against gold standard SNP6 microarray and WES datasets with matched normal samples. Our workflow further classifies SNVs by somatic status and then uses this information to infer somatic mutational signatures and tumor mutational burden (TMB).

**Results:** Application of our workflow to tumor-only WES data produces tumor purity and ploidy estimates that are highly concordant with estimates from SNP6 microarray data and matched-normal WES data. The presence of cancer type-specific somatic mutational signatures was inferred with high accuracy. We also demonstrate high concordance of TMB between our tumor-only workflow and matched normal pipelines.

**Conclusion:** The proposed workflow provides, to our knowledge, the only open-source option for comprehensive allele-specific CNA analysis and SNV classification of tumor-only WES with demonstrated high accuracy.

## Introduction

Copy number alterations (CNAs) are typically measured by the ratio of tumor to normal DNA abundance. However, tumor purity and ploidy affect this ratio and must be incorporated to infer absolute copy numbers (Carter et al., 2012; Van Loo et al., 2010). Information from heterozygous germline single nucleotide polymorphisms (SNPs) further allows deconvolution of absolute copy number into the two parental copy numbers. This parental or allele-specific copy number provides a direct readout of LOH (when either the maternal or paternal copy is lost), which can indicate the complete loss of wild-type function when a somatic mutation in a putative tumor suppressor is identified (Knudson, 1971). Inferring allele-specific copy number is further crucial to understanding mutagenesis, allowing determination of clonality and timing of copy number changes at the same locus (Carter et al., 2012; McGranahan et al., 2015; Nik-Zainal et al., 2012).

Whole exome sequencing (WES) and targeted panel sequencing become routine applications in the clinic, providing comprehensive data while saving cost and scarce tumor tissue by eliminating the need for multiple single analyte assays. Such comprehensive tests may therefore aid treatment decision-making by increasing the detection of actionable alterations, which includes point mutations and amplifications of oncogenes in targeted therapies, microsatellite instability (MSI), and tumor mutational burden (TMB) in immunotherapy (Chalmers et al., 2017; Zehir et al., 2017).

Sequencing both tumor and matched normal specimens currently provides certain benefits over tumor-only sequencing, even in diagnostic settings where alterations of uncertain significance are usually ignored. For example, high depth sequencing of blood samples can more reliably identify hotspot mutations that arose in heme rather than in tumor cells (Coombs et al., 2017; Steensma et al., 2015). Matched normal samples are also commonly required for existing algorithms to detect complex biomarkers such as MSI, TMB, or LOH. Obtaining comprehensive information from clinical tumor-only sequencing data could reduce time and cost, while enabling analyses of the large number of archived specimens for which blood samples are unavailable. However, the reliability of tumor-only sequencing is not well assessed (Jones et al., 2015; Shi et al., 2018).

Without matched normal samples, it is necessary to distinguish algorithmically between somatic mutations and germline variants. Existing approaches commonly involve machine learning using public germline and somatic databases, *in silico* predictions of the functional impact of mutations, as well as allelic fractions (the ratios of non-reference to total sequencing reads) of mutations and their neighboring SNPs (Kalatskaya et al., 2017; Smith et al., 2016). Recently developed tools for high coverage sequencing additionally use allele-specific copy number, allowing the calculation of accurate posterior probabilities for all possible somatic and germline genotypes (Halperin et al., 2017; Riester et al., 2016; Sun et al., 2018). However, complete and thoroughly benchmarked workflows are lacking.

We present a complete workflow for tumor-only hybrid-capture data, benchmark it against two gold standard datasets of matched normal WES sequencing and Affymetrix SNP6 microarrays, and compare it to alternative methods (Shen and Seshan, 2016). Using the ovarian carcinoma (OV) and lung adenocarcinoma (LUAD) datasets of The Cancer Genome Atlas (TCGA), which represent opposing extremes with respect to tumor purity, copy number heterogeneity and TMB, we demonstrate high reliability of tumor-only analyses for inference of allele-specific copy number, identification of functional mutations, LOH, mutational signatures, and TMB.

## Results

Reliable analysis of clinical tumor-only sequencing data involves multiple non-trivial steps that are distinct from the analysis of matched tumor and normal sequencing. Below we describe and benchmark a detailed workflow for hybrid-capture tumor-only sequencing data including variant calling, coverage normalization for copy number calling, purity and ploidy inference, and classification of variants by somatic status (Supplementary Fig. 1).

### Tumor purity and ploidy inference

We selected ovarian carcinoma (OV) and lung adenocarcinoma (LUAD) WES data from TCGA as complementary, representative datasets for our benchmarking study (Cancer Genome Atlas Research |Network, 2011, 2014). Among the TCGA datasets, OV shows the highest tumor purity due to the availability of large surgical specimens. High purity complicates somatic vs. germline classification due to the overlapping distributions of expected allelic fractions. The LUAD dataset, obtained by core needle biopsies, in contrast ranks among lowest in tumor purity, presenting a different challenge for copy number calling because of the dilution of signal (Riester et al., 2016; Sun et al., 2018). LUAD is additionally challenging because of increased copy number heterogeneity (Carter et al., 2012; Zack et al., 2013). Sub-clonal copy number changes increase the number of copy states, making ploidy inference often ambiguous (Carter et al., 2012; Ha et al., 2014).

We first compared maximum likelihood purity and ploidy estimates from our workflow using tumor WES with those from manually curated ABSOLUTE SNP6 microarray calls (Fig. 1A-D, Supplementary Table 1). We analyzed 233 OV and 442 LUAD samples, and found a high correlation of microarray and WES results for tumor purity (Pearson correlation of 0.75 and 0.84 for OV and LUAD, respectively) and tumor ploidy for OV (Pearson correlation of 0.73). Ploidy estimates for LUAD also showed a high concordance for the majority of samples (Pearson correlation of 0.57). Additionally, we applied FACETS, a widely used allele-specific CNA analysis tool for tumor and matched normal sequencing (Shen and Seshan, 2016), to both OV and LUAD paired WES data (Supplementary Fig. 2). For 68.9% of all samples, all 3 tools generated concordant purity and ploidy calls (Supplementary Fig. 3). For OV, PureCN showed a higher ploidy concordance with ABSOLUTE than FACETS (87.1% vs. 73.8%) whereas for LUAD, its concordance was slightly lower (77.1% vs. 79.6%).

**Figure 1.**
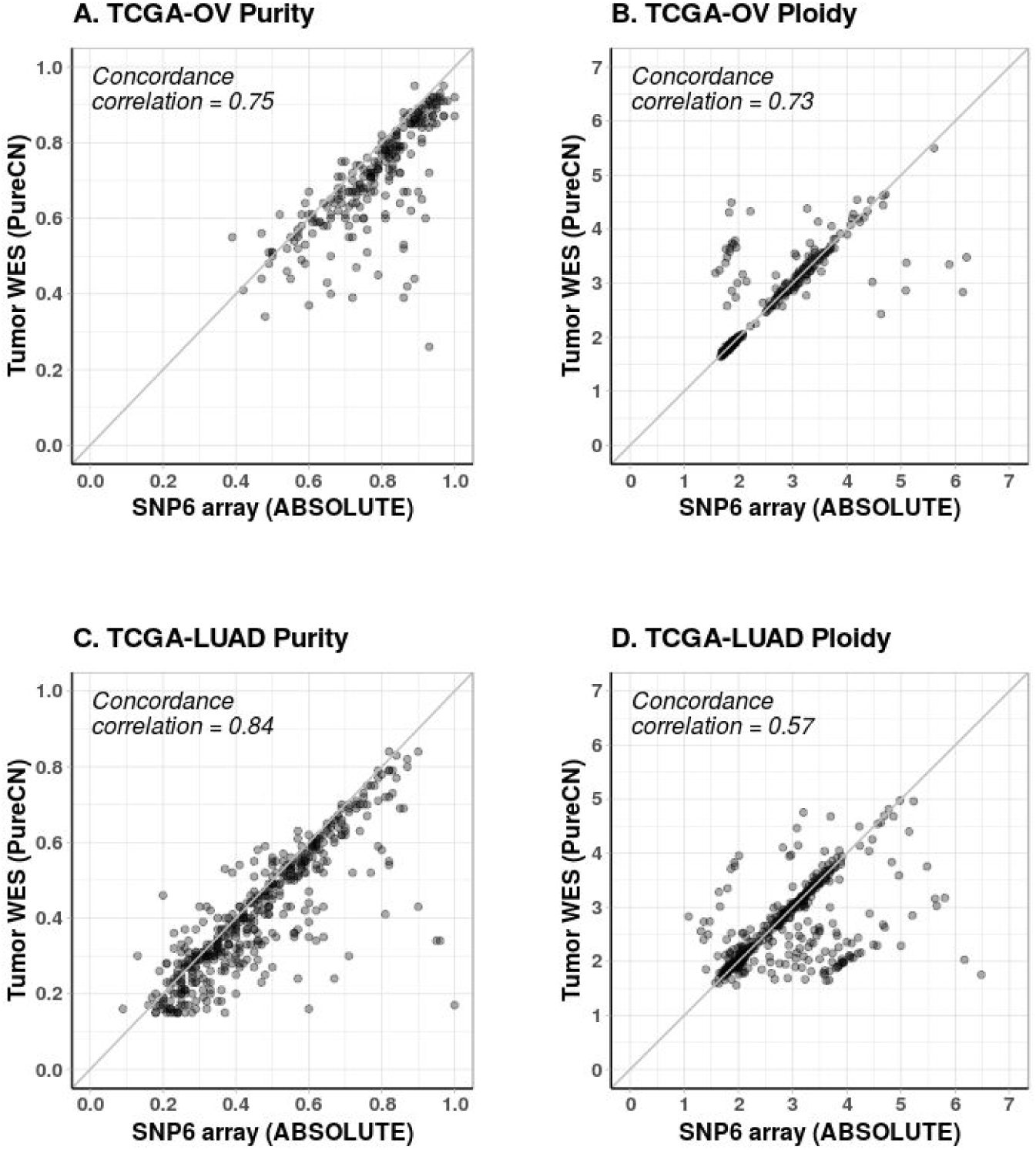
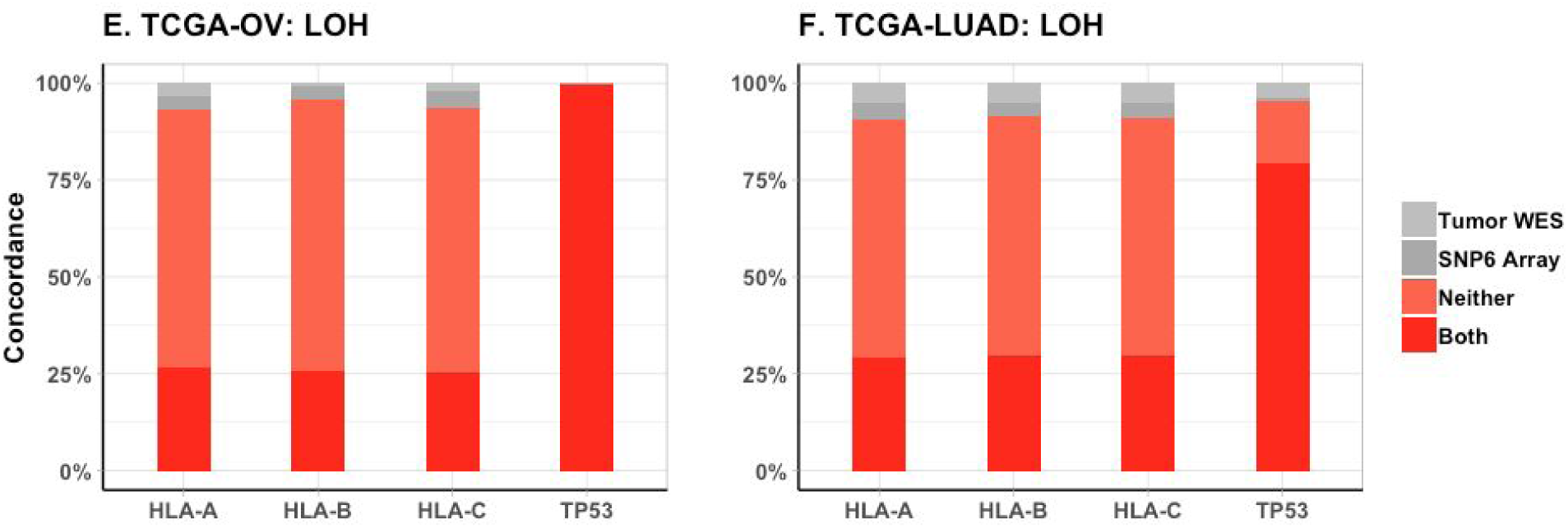
Accuracy of purity, ploidy and LOH inference. (A-D) Comparison of purity and ploidy estimates from paired SNP6 microarray data (ABSOLUTE, (Carter et al., 2012)) against those from tumor-only WES data (PureCN) in OV and LUAD samples. (E-F) Concordance of loss of heterozygosity (LOH) calls between the two analyses was reviewed on HLA-A/B/C and TP53 loci for the cases with sufficient power to detect LOH. LOH observed in both microarray and WES analyses (*Both*, red); absent in both analyses (*Neither*, orange); detected only from microarray data (*ABSOLUTE*, dark grey); detected only from WES data (*PureCN*, light grey).

### Loss of heterozygosity

We demonstrated the accuracy of allele-specific copy number analysis by examining loss of heterozygosity (LOH) at two loci of main clinical interest, HLA-A/B/C and TP53. HLA and TP53 loci were investigated in 143 and 223 OV cases, respectively, where both tumor-only WES and SNP6 array made LOH calls (Supplementary Table 2). For LUAD, the same comparison was done in 298 and 332 samples for HLA and TP53 loci, respectively. In OV, the mean agreement in LOH status between tumor-only WES and SNP6 microarray was 94.2% for HLA and 99.6% for TP53 (Fig. 1E). In LUAD, it was 91.0% for HLA and 95.5% for TP53 (Fig. 1F), with the discordant samples showing low purity (average of 30.9% vs. 43.3%, Mann–Whitney p-value < 0.0005).

### Classification of variants by somatic status

We next evaluated the somatic status predictions of variants not found in public germline databases. We first compared predictions against a simple model that utilizes only allelic fractions. This essentially compared the performance of commonly used *ad hoc* allelic fraction filters such as 0.4 against our model that adjusts allelic fractions for allele-specific copy number. We observed a significant improvement over this simple model in tumors with purity over 30% (Fig. 2A-B, Supplementary Table 3). Below 30% tumor purity, inclusion of copy number does not provide a benefit for classification due to the large difference in expected allelic fractions of germline and somatic variants.

**Figure 2.**
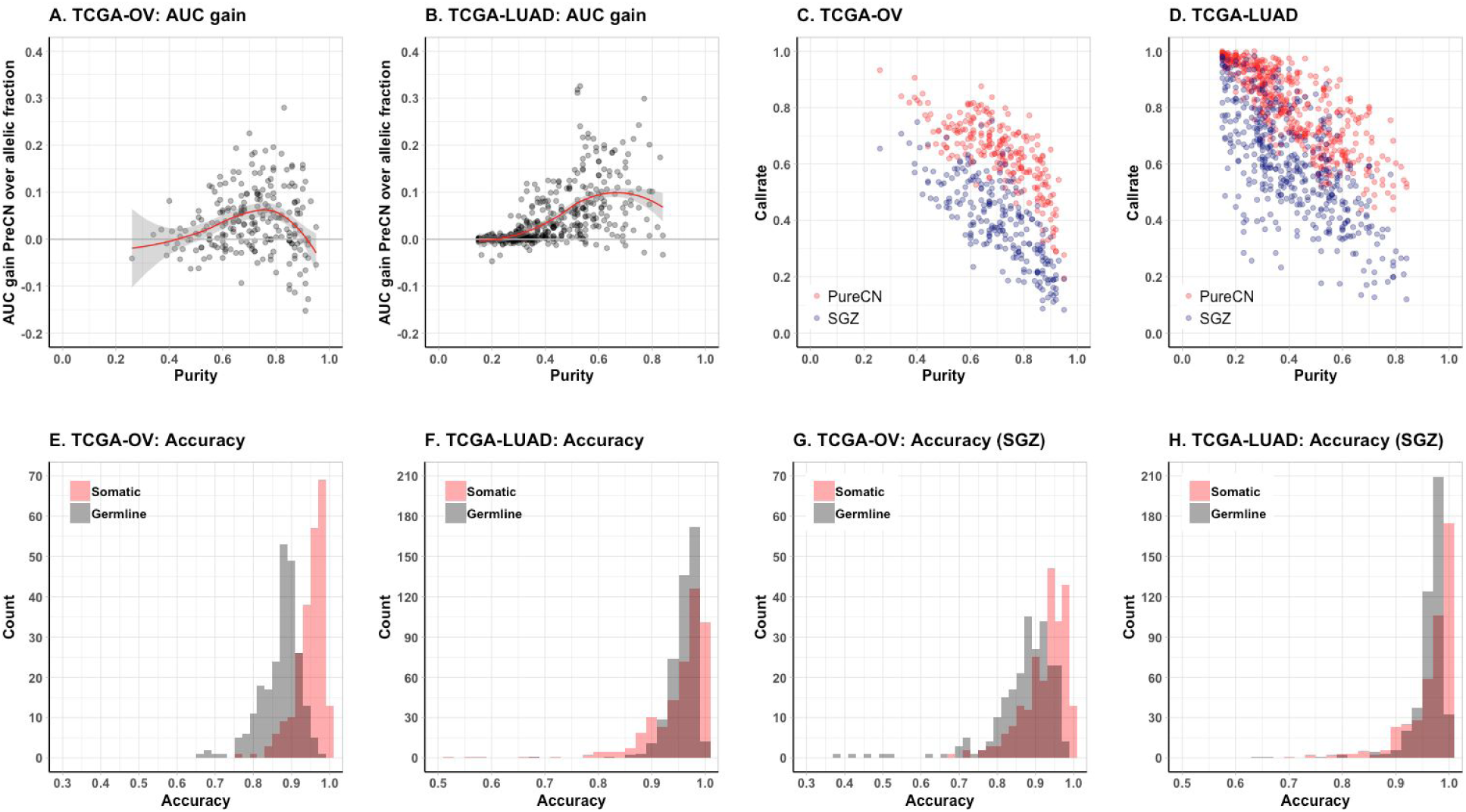
Accuracy of variant classification. (A-B) Gain in AUC of the somatic status prediction by PureCN over a model that only utilizes allelic fractions, shown as a function of tumor purity. (C-D) Correlation of tumor purity and call rates in OV and LUAD for PureCN (red) and SGZ (blue, Sun et al., 2018). (E-H) Histograms of accuracy rates for all samples. These are the fractions of variants correctly called as somatic (red) or germline (grey).

We then examined how many variants can be classified with reasonable certainty (Supplementary Table 4). As expected, this call rate was largely a function of tumor purity (Fig. 2C-D). Increasing sequencing coverage also increased these rates (Supplementary Fig. 4A-B). Somatic variants were classified with higher median accuracy than germline variants (96.1% vs 88.1%, respectively in OV; and 97.2% vs 96.6%, respectively in LUAD, Fig. 2E-F). This is also expected, since the somatic group includes sub-clonal mutations, which are usually easier to classify than mono-clonal mutations due to their lower allelic fractions and therefore higher allelic fraction difference compared to germline. We observed a similar median somatic and germline accuracy using SGZ (94.0% and 88.9% in OV; and 98.4% and 97.3% in LUAD; Fig. 2G-H; (Sun et al., 2018)), but with lower median call rates (39.5% and 59.5% for OV and LUAC, respectively for SGZ vs. 64.4% and 82.2% for PureCN).

### Tumor mutational burden

We next sought to investigate the accuracy of the variant classification for determining tumor mutational burden (TMB). From the comparison of tumor-only and paired analysis modes, we found a high concordance (Pearson correlation 0.98) and good calibration of somatic mutation rates per megabase, in both OV and LUAD (Fig. 3A).

**Figure 3.**
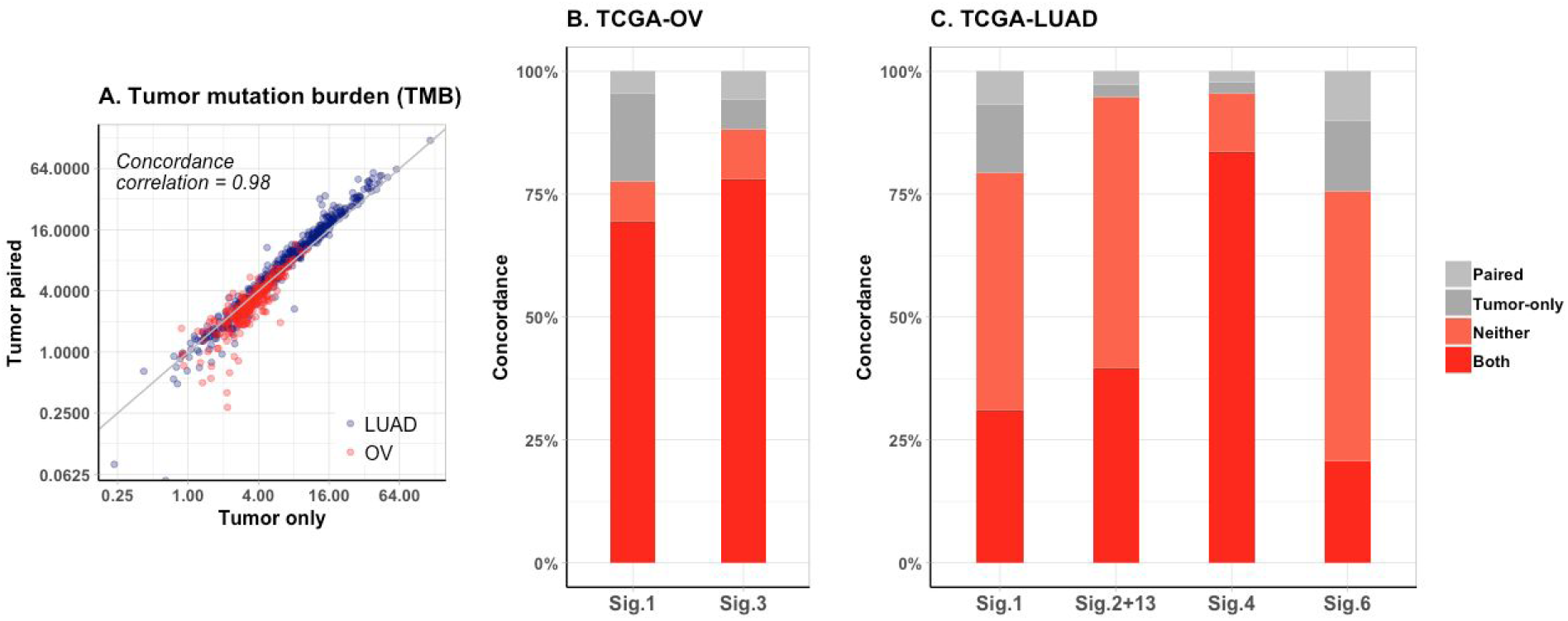
TMB and mutational signatures. (A) TMB from OV (red) and LUAD (blue) samples in tumor-only vs. paired modes shown on a log scale. (B-C) Concordance of COSMIC mutational signatures between tumor-only and paired analysis modes. Mutational signatures observed in both tumor-only and paired modes of analysis (*Both*, red); absent in both analyses (*Neither*, orange); detected only from tumor-only analysis (*Tumor-only*, dark grey); detected only from paired mode of analysis (*Paired*, light grey).

### Mutational signatures

To further evaluate the clinical utility of our workflow, we assessed the accuracy of mutational signature identification (Alexandrov et al., 2013) from tumor WES data with or without matched-normal profile. Among 30 validated mutational signatures, we investigated the two OV-associated mutational signatures with known aetiology in detail (Fig. 3B). Signature 1, generated by spontaneous deamination of 5-methylcytosine, has been found in all cancer types and is linked to aging. Signature 3 is associated with homologous repair deficiency, a potential biomarker for PARP inhibition in ovarian cancer (Moore et al., 2018). We obtained a high agreement for mutational signature calls from tumor-only and paired analyses (77.5% for Signature 1 and 88.1% for Signature 3), confirming that our workflow can reliably detect mutational signatures without matched normal profile even in high purity samples.

In LUAD data we again reproduced previously associated signatures of known aetiology. In addition to the aging Signature 1, we found a significant fraction of samples dominated by the signatures APOBEC (Signature 2 and Signature 13), tobacco (Signature 4), and DNA mismatch repair deficiency (Signature 6) signatures (Fig. 3C, Supplementary Table 5). We observed high agreement between tumor-only with matched normal data for all these signatures (79.3%, 94.8%, 95.7%, and 75.5% for Signatures 1, combined 2+13, 4 and 6, respectively).

## Discussion

We present an easy-to-implement workflow for reliable analysis of clinical tumor-only whole exome sequencing data without matched normal samples. This workflow is validated on OV and LUAD data from TCGA and benchmarked against a gold-standard, manually-curated analysis of SNP6 microarray data with matched normals. Our workflow estimates tumor purity, ploidy, LOH, TMB and mutational signatures with a high concordance to established workflows for SNP6 and whole-exome sequencing data with paired tumor and normal samples.

To our knowledge, this is the first thoroughly validated open source tumor-only TMB pipeline. TMB is an emerging biomarker for response to immunotherapy (Goodman et al., 2017; Rizvi et al., 2015; Rosenberg et al., 2016; Snyder et al., 2014), but the current lack of standards significantly challenges implementing TMB testing in the clinic (Büttner et al., 2019). We believe this open source reference implementation will help establish standards for TMB calling.

While high tumor purity challenges somatic status classification (Fig. 2C-D), the proposed approach to determining clinically relevant biomarkers such as TMB and somatic signatures is surprisingly robust to varying tumor purity (Fig. 3). Notably, signatures of clear aetiology such as homologous repair deficiency, APOBEC, or smoking had a significantly higher concordance with matched analyses than broader and less certain signatures, for example those associated with aging. In contrast, we also note that high tumor purity is beneficial for LOH and copy number calling. Still, all parts of the workflow achieved high accuracy in tumors of 40-60% purity, the range in which most clinical tissue specimens fall.

This study has several limitations. First, we focused on benchmarking our tumor-only workflow where it differs from standard matched tumor and normal analyses. A systematic evaluation of accuracy for the variant calling steps upstream of this workflow is beyond the scope of this study (Krøigård et al., 2016; Xu, 2018). Second, our workflow is currently not designed for whole-genome sequencing (WGS) data. In contrast to gold-standard WGS tools, PureCN was designed for high coverage data (>100X) and currently does not use information largely unavailable in hybrid-capture data such as split reads or SNP phasing. These would be straightforward additions once high coverage diagnostic WGS becomes common in oncology. Third, as with allele-specific CNA calling in matched tumor and normal data, purity and ploidy inference can be ambiguous in a minority of cases of low purity or of high heterogeneity. Our pipeline therefore provides tools that allow manual correction of results by trained curators, described in the documentation of the PureCN package. Importantly, the accuracy of TMB calling was robust even to inaccuracies in ploidy, partly because different ploidy solution can be equivalent for variant classification (Sun et al., 2018). Lastly, reliable labelling of clonal hematopoiesis from tumor-only or low coverage matched normal sequencing remains a shortcoming, but is an area of research we are currently pursuing.

Due to the high concordance with matched tumor and normal sequencing, the proposed workflow supports the clinical application of tumor-only sequencing, especially in diagnostic settings.

## Methods

### Installation

PureCN can be obtained and installed under the Artistic 2.0 (https://doi.org/doi:10.18129/B9.bioc.PureCN), (https://bioconda.github.io/recipes/bioconductor-purecn/README.html) license from or Bioconductor Bioconda GitHub (https://github.com/lima1/PureCN). Unless otherwise specified, .R scripts referred to below are part of the PureCN package.

### Dependencies

Genome reference FASTA files were downloaded from NCBI (ftp://ftp.ncbi.nlm.nih.gov/genomes/all/GCA/000/001/405/GCA_000001405.15_GRCh38/seqs_for_alignm ent_pipelines.ucsc_ids/GCA_000001405.15_GRCh38_no_alt_analysis_set.fna.gz), dbSNP version 135 was downloaded from https://software.broadinstitute.org/gatk/download/bundle and COSMIC version 77 from https://cancer.sanger.ac.uk/cosmic.

The workflow described in this paper assumes that germline SNPs and somatic mutations were identified by MuTect 1.1.7 (Cibulskis et al., 2013), which requires Java 1.7. Other commonly used variant callers with tumor-only mode can be used instead, but the resulting VCFs need to be filtered for common artifacts before subjecting them to this workflow.

This workflow further requires a read mappability file for the reference FASTA file. The mappability score *m_i_* is defined as 1/(# alignments) for a k-mer starting at position *i* in the reference genome. For hg19, these scores can be downloaded as precomputed for various k-mer sizes from the UCSC genome browser (http://hgdownload.cse.ucsc.edu/goldenpath/hg19/encodeDCC/wgEncodeMapability/). For other reference genomes, we require the GEM library (Derrien et al., 2012) version 1.315.

Finally, GATK3 `CallableLoci` was used to collect callable regions with sufficient coverage, mappability, and sequence quality. VCF were combined into a single multi-sample VCF with GATK3 `CombineVariants`.

### Data download

Tumor and normal BAM files were downloaded through the GDC Data Transfer Tool using manifest files built by the `GenomicDataCommons` R/Bioconductor package (Morgan and Davis, 2017). The `TCGAutils` R/Bioconductor package (Ramos et al., 2018) was used to annotate the manifest file: `TCGAutils::UUIDtoBarcode` for transferring Universally Unique Identifiers (UUID) to TCGA barcodes and `TCGAutils::TCGAbiospec` for extracting biospecimen data from TCGA barcodes. Capture kit information related to each BAM file was obtained via the GDC API using `Curl`. BAM files mapping to multiple capture kits were excluded. BED files containing the locations of baits based on hg19 were lifted over to GRCh38 using `hg19ToHg38` liftover chain file downloaded from the UCSC Genome Browser (http://hgdownload.soe.ucsc.edu/goldenPath/hg19/liftOver/hg19ToHg38.over.chain.gz).

ABSOLUTE analysis of TCGA SNP6 microarray data has been described previously (Carter et al., 2012; Taylor et al., 2018; Zack et al., 2013). The manually curated ABSOLUTE output was obtained from Synapse (https://www.synapse.org/#!Synapse:syn7416143) and lifted over to GRCh38.

None of the data analyzed in this study were used to develop or tune the algorithm or parameters and thus represent true validation sets.

### Variant calling

`MuTect` was run separately on both tumor and normal BAM files (Supplementary Fig. 1, purple and blue lines, respectively) using arguments `--dbSNP Homo_sapiens_assembly38.dbsnp.vcf` and `--cosmic CosmicCodingMuts.vcf`. For benchmarking purposes, MuTect was also run in matched normal mode by providing both tumor and normal BAM files and otherwise identical parameters (Supplementary Fig. 1, purple dashed line). Capture kit intervals were not provided to include all SNPs with sufficient coverage in the flanking regions of baits. Normal samples were run in artifact detection mode (`--artifact_detection_mode` argument) and then combined into a single multi-sample VCF (referred as `normal.panel.vcf.file` in the following) using GATK3 `CombineVariants` with argument `--minimumN` 5. The latter specifies the minimum number of normal VCF files containing the variant call to be included in the normal database. This was set to a high value to ensure that all individual-specific germline variants were ignored, which would otherwise indirectly provide matched normal information for some tumors.

### Reference files generation

The first step of the workflow (Supplementary Fig. 1) is the generation of reference files for each capture kit.

To exclude regions of low read mappability, bigWig files were generated from the GRCh38 reference genome assembly without ALT contigs using the GEM library (Derrien et al., 2012) and the UCSC `wigToBigWig` tool. The k-mer size in `gem-mappability` was set to the read lengths of the studies and the maximum number of mismatches and edit distances was set to 2 (`-m` and `-e` arguments, respectively), matching the settings used by ENCODE.

Next, the script `IntervalFile.R` was used with default arguments to annotate the regions defined by the baits BED file with mean GC-content, mean mappability, and gene symbols. `IntervalFile.R` further splits on- and off-target regions into bins of maximum 400 bp and 200 kbp, respectively (Supplementary Fig. 1, black line), as previously described (Talevich et al., 2016).

Only capture kits that were used for more than 100 samples were considered and separate normal databases were built for each kit (Supplementary Fig. 1, blue lines). With these criteria, 233 OV tumors and their matched normal samples were processed from two different capture kits (‘Custom V2 Exome Bait, 48 RXN X 16 tubes’ and ‘SureSelect Human All Exon 38 Mb v2’). For LUAD, 442 tumors and matched normal samples were processed (‘Custom V2 Exome Bait, 48 RXN X 16 tubes’ kit).

Coverage files were generated for all normal samples using `Coverage.R` (Supplementary Fig. 1, blue lines) with default arguments. This script normalizes on- and off-target coverages independently for GC-content.

The normal coverage databases for the LUAD and the two OV capture kits (output file `normalDB.rds`) were then generated with the `NormalDB.R` script with default arguments. In brief, outlier normal samples with very high or very low coverage were excluded (> 4x or < 0.25x coverage median, respectively). Furthermore, intervals with no read count in more than 3% of samples and average coverage lower than 25% of the chromosome median were removed. In total, 157 and 176 process-matched normals from two OV capture kits, and 250 from LUAD were used to build the three normal databases.

For each interval, `NormalDB.R` then calculates the inverse of the log2-copy number ratio standard deviation across all normal samples and creates the `interval_weights.txt` output file, later used by the segmentation function to downweight intervals with high variance in normal controls.

Reads harboring non-reference alleles have a lower chance of passing filters, thus resulting in average allelic fractions of heterozygous SNPs below the expected 0.5. Therefore `NormalDB.R` next computes a position-specific non-reference mapping bias (output file `mapping_bias.rds`) for all variants in the `normal.panel.vcf.file`, provided through the `--normal_panel` argument. Mapping bias is defined as the ratio of the sum of all alt reads over all samples vs. the total number of reads of heterozygous SNPs (allelic fraction > 0.05 and < 0.9). This procedure further uses an empirical Bayes approach that adds the average number of non-reference and total reads per SNP across all samples to this ratio, thus forcing the mapping bias of rare or low coverage SNPs closer to the average mapping bias.

### Whole exome copy number calling

Tumor coverages were calculated and GC-normalized using the `Coverage.R` script with default arguments, analogous to the normal coverages (Supplementary Fig. 1, red line).

The `PureCN.R` script was then used for the main copy number calling step that includes tumor purity and ploidy inference as well as classification of somatic status and clonality for all variants (Riester et al., 2016). The `--postoptimize` flag as well as all previously mentioned reference files were provided. Variants in the UCSC simple repeat track were excluded (`--snpblacklist` argument). Otherwise default parameters were used.

In brief, tumor vs. normal log2-copy number ratio was first calculated and denoised using tangent normalization (Beroukhim et al., 2010), again independently for on- and off-target regions. Mapping bias of variants not found in the normal database was imputed by averaging the mapping bias of the 5 neighbors on both sides, weighting each of the 10 SNPs by corresponding number of samples in the database. Then DNAcopy (Venkatraman and Olshen, 2007) was used for the segmentation of merged on- and off-target log2-ratios. Reliable germline SNPs present in the normal database without major mapping bias were used to improve the segmentation by Ward clustering and identification of copy-neutral LOH. Candidate purity and ploidy combinations for the segmented log2-ratios were identified in a 2D-grid search, and subsequently optimized using Simulated Annealing. Allelic variants were finally fitted to all local optima, calculating somatic posterior probabilities for all variants.

The likelihood model of PureCN has been described previously (Riester et al., 2016). PureCN versions > 1.8.0 differ in two minor details. First, the uncertainty of copy number log2-ratio standard deviation is now included in the optimization. This is an advantage in high quality samples where shifts in log2-ratio across chromosomes can sometimes exceed the average noise within segments. Second, the observed sample ploidy can differ from the true ploidy, especially in smaller gene panels that cover only small fractions of the genome. Previously described PureCN versions modeled this potential deviation as a function of the sample noise, but this was changed to a function of tumor purity.

FACETS version 0.5.6 was used (Shen and Seshan, 2016). Tumor and normal bam file pairs were processed by snp-pileup with the parameters `-g -q15 -Q20 -P100 -r25,0`, and the outputs from which were imported using `readSnpMatrix` and further processed by `preProcSample`, `procSample` with `cval = 150`, and `emcncf`.

### Classification of variants by somatic status

Variants with a somatic posterior probability ≥ 0.8 were classified as somatic, ≤ 0.2 as germline. While this cutoff may seem arbitrary and liberal, the assumption is that such a classification of specific variants is mostly of interest when additional information strongly suggests functional significance, such as determined by *in silico* functional prediction tools or due to location in hotspot domains of relevant genes. All variants found in germline databases with small prior probability of being somatic were excluded from benchmarking.

SGZ (Sun et al., 2018) in version 1.0.0 was applied to all WES data. SGZ is methodologically similar to PureCN, but does not include the uncertainty of allele-specific copy number in the posterior probability calculation and is not correcting allelic fractions for non-reference mapping bias. Since SGZ does not ship with a copy number tool, allele-specific copy number data as generated by the PureCN `callLOH` function was provided. Variants flagged by PureCN for recurrent presence in the pool of normals or for high imputed mapping bias were excluded. The same set of variants was thus used for both tools. Variants labeled “germline”, “probable germline”, “somatic”, “probable somatic” or “somatic subclonal” by SGZ were considered called, and all others uncalled. Parameters of both tools including classification cutoffs were specified before data analysis.

### Tumor mutational burden

To call TMB, defined as the number of somatic mutations per megabase, `Dx.R` was run with the `--callable` and `--snpblacklist` flags and otherwise default arguments, confining the regions of interest to bases reliably callable by MuTect and excluding simple repeats. Callable regions were obtained by GATK3 `CallableLoci` with a minimum read depth of 30 (`--minDepth` argument) and otherwise default parameters. Non-coding regions were excluded from the `CallableLoci` output using `FilterCallableLoci.R`. Mutations with a posterior probability above 0.5 for being somatic, which were also not included in germline databases and not flagged by PureCN, were included in the TMB calculation. In the matched tumor and normal TMB pipeline, somatic variants were assigned a prior somatic probability of 0.999 and germline SNPs a prior of 0.0001; otherwise identical parameters were used.

### Mutational signatures

To identify the 30 mutational signatures (Alexandrov et al., 2015) curated by the Wellcome Trust Sanger Institute (http://cancer.sanger.ac.uk/cosmic/signatures), `Dx.R` was run with the non-coding regions were kept to increase the number of mutations. Samples with less than or equal to 50 somatic mutations were excluded as recommended in (Rosenthal et al., 2016), leaving 160 OV and 368 LUAD samples for analysis.

### Statement of Reproducible Research

Analyses presented in this manuscript are reproducible using the code available through https://github.com/shbrief/CNVWorkflow_Code.

## Supporting information

Supplementary Table 1. Purity and ploidy estimates from ABSOLUTE, PureCN (tumor-only and paired mode), and FACETS.

Supplementary Table 2. LOH of HLA and TP53 loci from ABSOLUTE and PureCN.

Supplementary Table 3. AUC gain of PureCN in 223 OV and 441 LUAD samples.

Supplementary Table 4.1. Benchmark results using PureCN.

Supplementary Table 4.2. Benchmark results using SGZ

Supplementary Table 5. COSMIC mutational signatures of OV and LUAD samples in tumor-only and paired analysis modes.

## Supplementary Figures

**Supplementary Figure 1.**
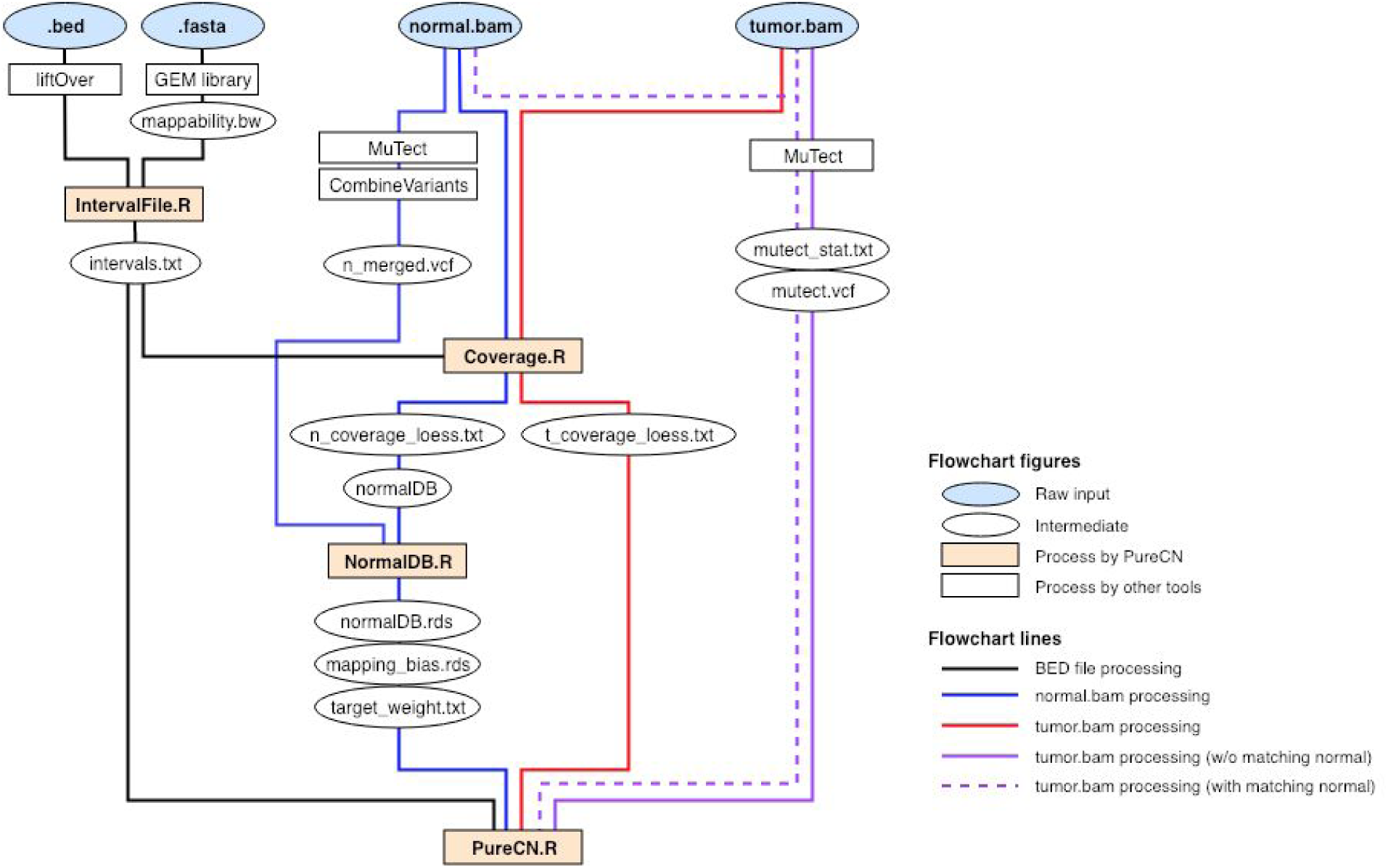
CNA analysis workflow. Raw input data files and the intermediate/processed data files are labeled with blue and white oval shapes, respectively. R scripts provided by PureCN are depicted by orange squares, and third party tools by white squares. Black solid lines indicate how the target region information is processed. Blue and Red solid lines describe how normal and tumor bam files are processed, respectively. Dashed and solid purple lines are showing how germline SNPs and somatic mutations were prepared with or without matched normal, respectively.

**Supplementary Figure 2.**
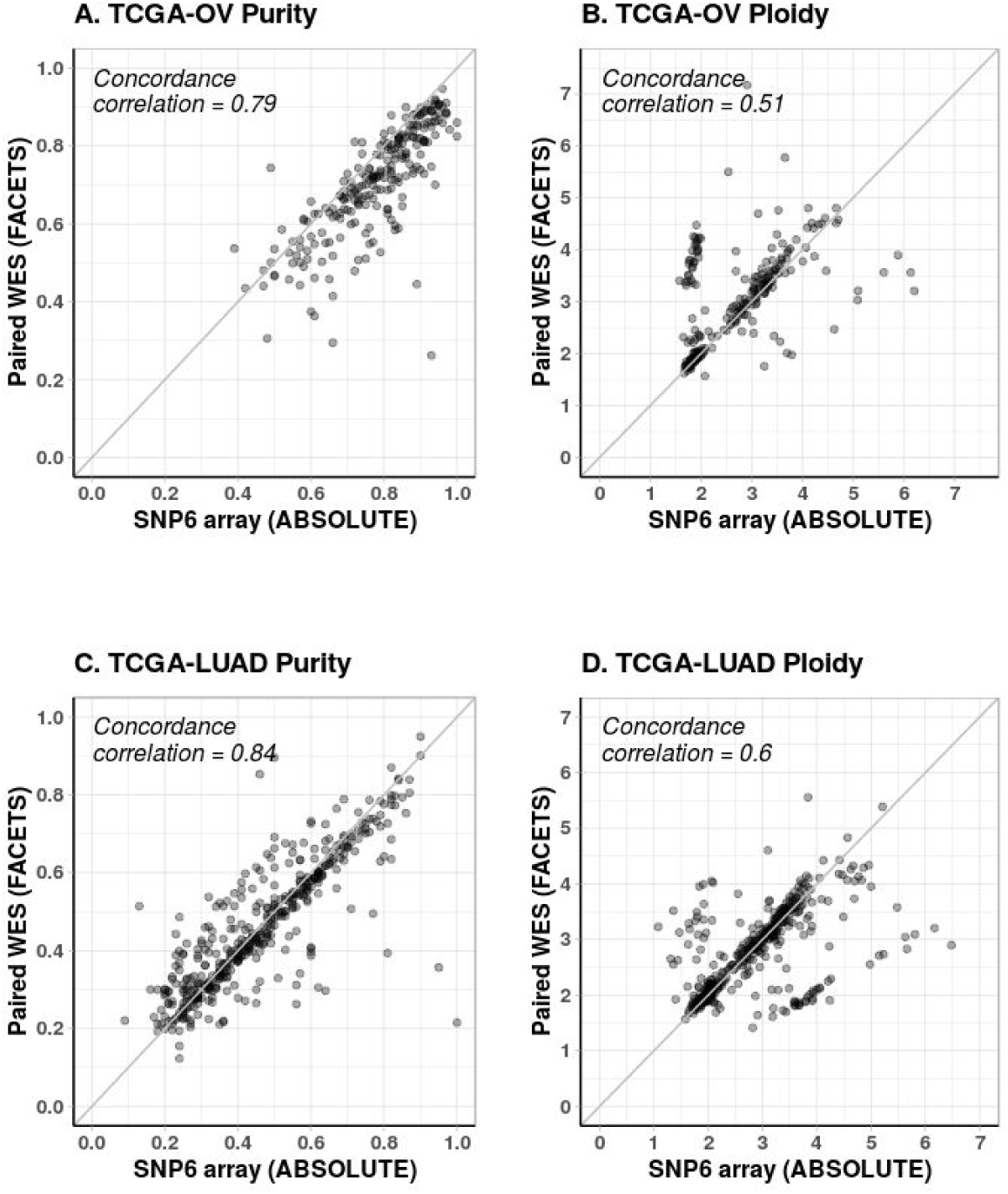
Purity and ploidy estimates using an alternative tool. Purity and ploidy estimates from paired WES data were obtained using FACETS. As in Fig. 1, 233 OV and 442 LUAD samples were analyzed and compared to ABSOLUTE calls. (A) and (B) are purity and ploidy estimates of OV, respectively. (C) and (D) are purity and ploidy estimates of LUAD, respectively. For panel (C), 436 cases are plotted as FACETS did not return a purity estimate for 6 of the LUAD samples due to insufficient information.

**Supplementary Figure 3.**
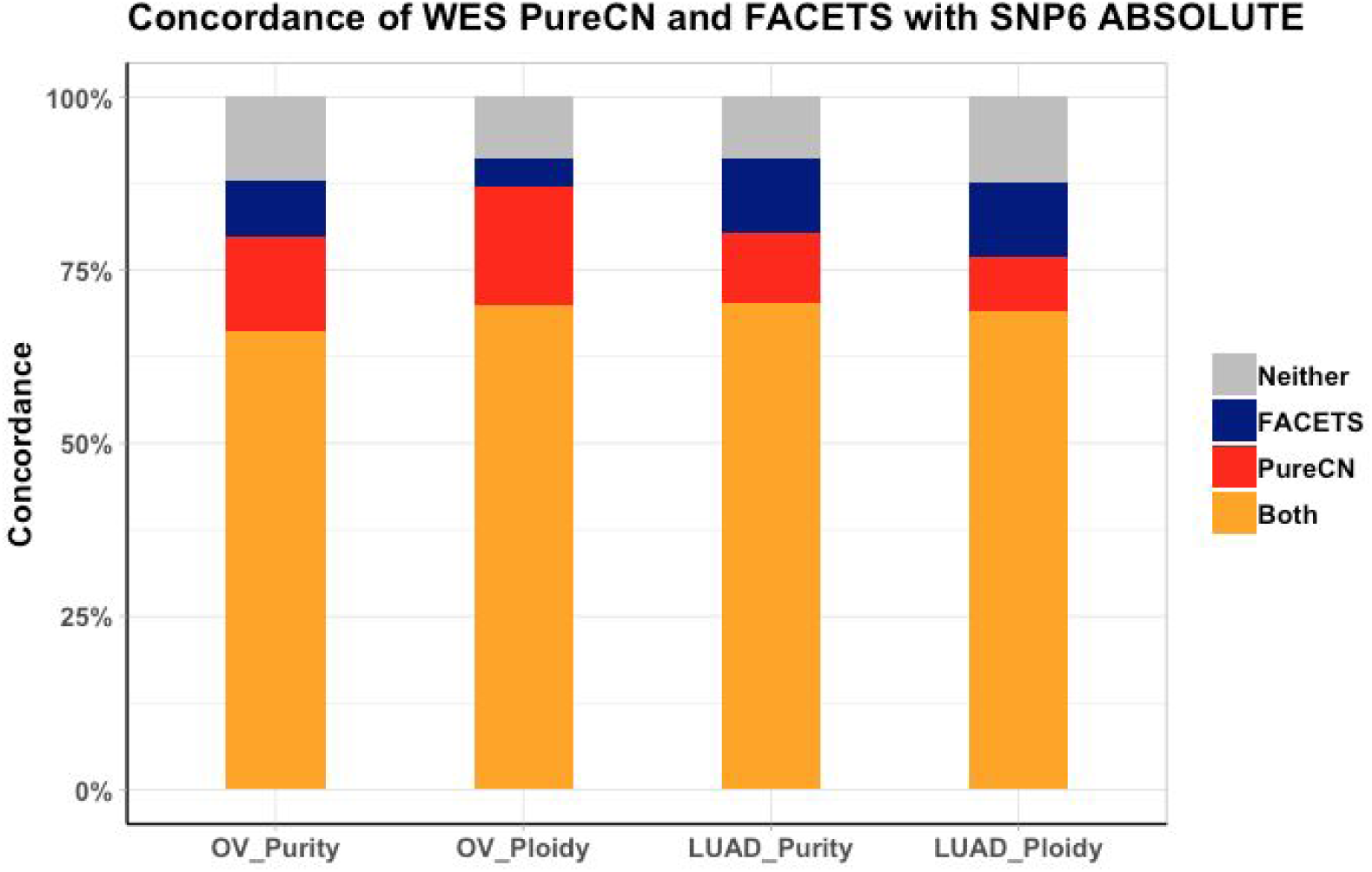
Concordance of PureCN and FACETS with ABSOLUTE. From 233 OV and 436 LUAD cases, concordance was calculated of WES-based estimates from PureCN and FACETS with SNP6 array-based ABSOLUTE calls. Concordance was defined as purity difference < 0.1 and a ploidy difference < 0.5. Estimates agreed by all three methods (*Both*, yellow); agreed by ABSOLUTE and PureCN only (*PureCN*, red); agreed by ABSOLUTE and FACETS only (*FACETS*, blue); neither PureCN nor FACETS agree with ABSOLUTE (*Neither*, light grey).

**Supplementary Figure 4.**
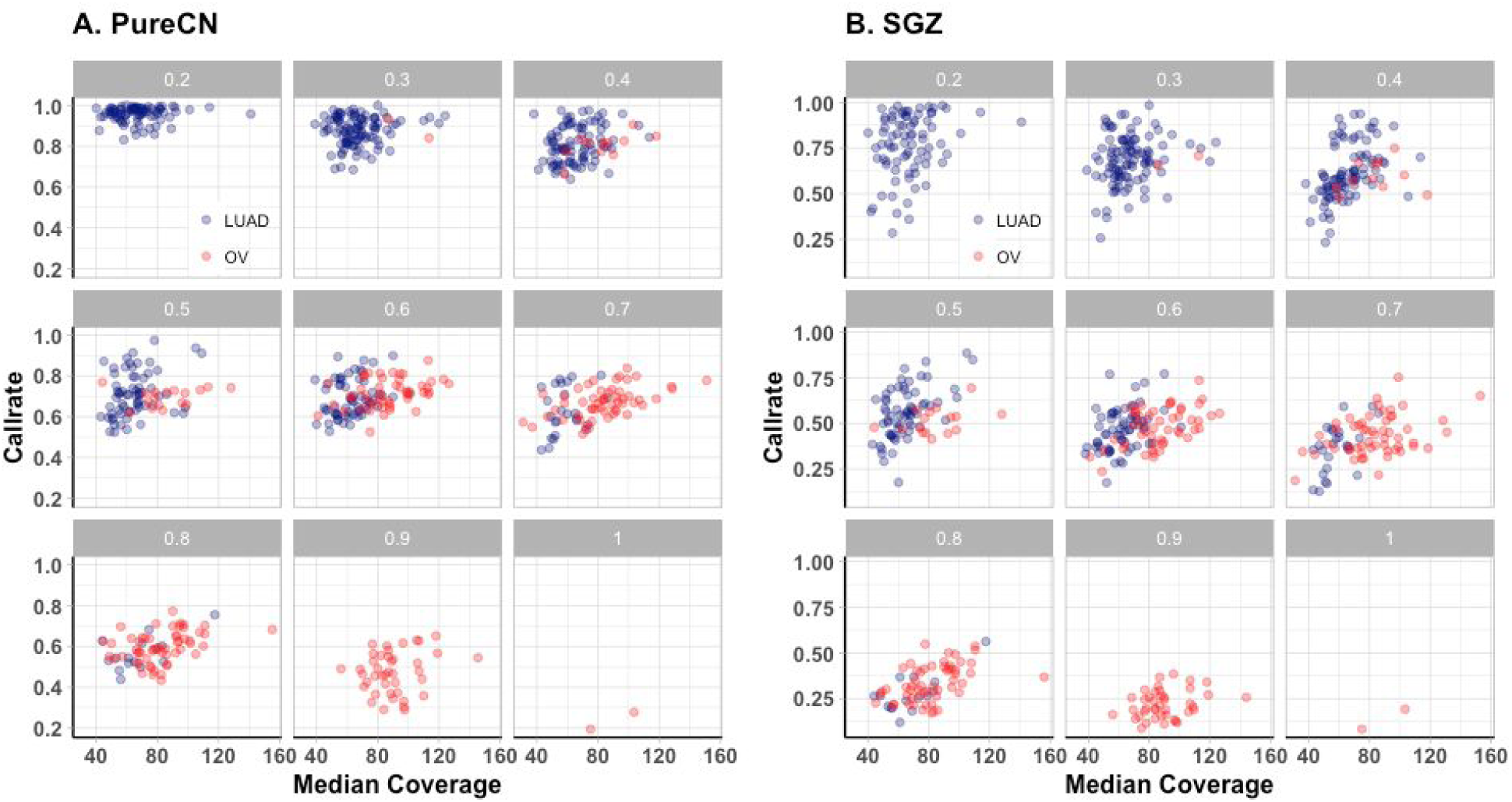
Correlation of call rates and median sequencing coverage. Median coverage is plotted against call rate for different purity ranges.

## Supplementary Tables

**Supplementary Table 1**. Purity and ploidy estimates from ABSOLUTE, PureCN (tumor-only and paired mode), and FACETS.

**Supplementary Table 2**. LOH of HLA and TP53 loci from ABSOLUTE and PureCN.

**Supplementary Table 3**. AUC gain of PureCN in 223 OV and 441 LUAD samples.

**Supplementary Table 4**. Benchmark results using PureCN (sheet_1) and SGZ (sheet_2).

**Supplementary Table 5**. COSMIC mutational signatures of OV and LUAD samples in tumor-only and paired analysis modes.

